# Breeding of a near-isogenic wheat line resistant to wheat blast at both seedling and heading stages through incorporation of *Rmg8*

**DOI:** 10.1101/2023.07.12.546477

**Authors:** Motohiro Yoshioka, Mai Shibata, Kohei Morita, Masaya Fujita, Koichi Hatta, Makoto Tougou, Yukio Tosa, Soichiro Asuke

## Abstract

Wheat blast caused by *Pyricularia oryzae Triticum* pathotype (MoT) has been transmitted from South America to Bangladesh and Zambia and is now spreading in these countries. To prepare against its further spread to Asian countries, we introduced *Rmg8*, a gene for resistance to wheat blast, into a Japanese elite cultivar, Chikugoizumi (ChI), through recurrent backcrosses, and established ChI near-isogenic lines, #2-1-10 with the *Rmg8*/*Rmg8* genotype and #4-2-10 with the *rmg8*/*rmg8* genotype. A molecular analysis suggested that at least 96.6% of the #2-1-10 genome was derived from the recurrent parent ChI. The #2-1-10 line was resistant to MoT not only in primary leaves at the seedling stage but also in spikes and flag leaves at the heading stage. The strength of the resistance in spikes of this *Rmg8* carrier was comparable to that of a carrier of the 2NS segment which has been the only one genetic resource released to farmer’s field for wheat blast resistance. On the other hand, the 2NS resistance was not expressed on leaves at the seedling stage nor flag leaves at the heading stage. Considering that leaf blast has been increasingly reported and regarded as an important inoculum source for spike blast, *Rmg8* expressed at both the seedling and heading stages, or more strictly in both leaves and spikes, is suggested to be useful to prevent the spread of MoT in Asia and Africa.

## INTRODUCTION

Wheat production in the world is currently threatened by the spread of the blast disease caused by *Pyricularia oryzae* (syn. *Magnaporthe oryzae*) *Triticum* pathotype (MoT, the wheat blast fungus) (Cruz and Valent 2017). This pathogen first emerged in Brazil in 1985, then spread to its neighbor countries, i.e., Bolivia in 1996, Paraguay in 2002, and Argentina in 2007 (Singh et al. 2021; Valent et al. 2021). Recently, MoT further spread to Bangladesh (in 2016) and Zambia (in 2018) probably through infected seeds, and caused severe outbreaks of wheat blast in these countries (Islam et al. 2016; Malaker et al. 2016; Tembo et al. 2020). Molecular analyses suggested that these outbreaks were caused by MoT strains that were closely related to the South American strain B71, and were transmitted from South America to Asia and to Africa through independent introductions (Latorre et al. 2023; Liu et al. 2022).

To mitigate this devastating disease, resistance genes are needed. Intensive screening of common wheat and its relatives in Brazil, Bolivia, and Paraguay resulted in identification of several varieties with moderate resistance, many of which had the CIMMYT genotype ‘Milan’ in their pedigree (Singh et al. 2021). This resistance was proved to be conferred by a 2NS chromosomal segment (Cruz et al. 2016) which had been introduced from *Aegilops ventricosa* through translocation (2NS/2AS translocation) (Helguera et al. 2003). A newly developed variety carrying this 2NS segment, ‘BARI Gom33’, showed sufficient resistance to wheat blast in field tests in Bolivia and Bangladesh, and was released in Bangladesh in 2017 (Hossain et al. 2019). However, new MoT strains overcoming the 2NS resistance have already emerged in South America (Cruppe et al. 2020; Cruz et al. 2016), indicating an urgent need of novel resistance sources. Although intensive screenings were performed with many wheat lines from CIMMYT, South Asia, China, South America, etc. (Juliana et al. 2020; He et al. 2021; Phuke et al. 2022; Roy et al. 2021; Wu et al. 2022), the common finding was that the 2NS translocation provided the only major and consistent resistance source (Singh et al. 2021).

Resistance genes against wheat blast must fulfill at least two requirements so as to be useful in farmer’s fields. First, they must be effective at high temperature because this disease is severe at high temperature with an optimum between 25 and 30°C (Valent et al. 2021). Second, they must be effective in spikes because spike blast is the predominant form of this disease in the field (Valent et al. 2021). Tosa and his coworkers have identified five resistance genes against MoT, i.e., *Rmg2*, *Rmg3*, *Rmg7*, *Rmg8*, and *RmgGR119* (Anh et al. 2015; Tagle et al. 2015; Wang et al. 2018; Zhan et al. 2008). *Rmg2* and *Rmg3* identified in common whet cultivar ‘Thatcher’ (Zhan et al. 2008) were not effective at high temperature or in spikes. *Rmg7* identified in emmer wheat was effective in spikes (Tagle et al. 2015) but not at high temperature (Anh et al. 2018). On the other hand, *Rmg8* identified in common wheat cultivar ‘S-615’ was effective in spikes (Anh et al. 2015) and at high temperature (Anh et al. 2018). *Rmg8* was located on a distal region of the long arm of chromosome 2B, which was syntenic to the region of chromosome 2A harboring *Rmg7* (Anh et al. 2015). Furthermore, *Rmg8* and *Rmg7* recognized the same avirulence gene, *AVR-Rmg8* (Anh et al. 2018). These results suggested that *Rmg8* and *Rmg7* might be homoeologous genes derived from a single ancestral gene (Anh et al. 2018). *Rmg8* was present in ∼4% of local landraces of common wheat in the world (Wang et al. 2018). One of these *Rmg8* carriers harbored an additional resistance gene tentatively designated as *RmgGR119* (Wang et al. 2018).

It is important to monitor the distribution of avirulence genes in MoT populations to identify the most appropriate resistance genes to be used in breeding programs (Navia-Urrutia et al. 2022). *AVR-Rmg8* in MoT isolates is composed of several variants, e.g., eI, eII, eII’ (Wang et al. 2018), eII’’ (var4 in Navia-Urrutia et al. 2022), and eII’’’ types (Latorre et al. 2023). The eI type is strongly recognized by *Rmg8*, but the other types partially or fully evade the recognition by *Rmg8* (Holo et al. 2020; Latorre et al. 2023). Unfortunately, the latter (virulent or moderately virulent) types are already distributed widely in South America (Horo et al. 2020; Navia-Urrutia et al. 2022), suggesting that *Rmg8* is not useful in this continent where MoT originated (Navia-Urrutia et al. 2022; Valent et al. 2021). On the other hand, all isolates collected in Bangladesh and Zambia harbored the eI type and were actually avirulent on *Rmg8* carriers (Holo et al. 2020; Latorre et al. 2023). This result suggests that *Rmg8* is effective against the pandemic wheat blast population (Latorre et al. 2023), and will be useful in breeding for wheat blast resistance in Asia and Africa.

MoT is gradually spreading to northern districts in Bangladesh year after year (Singh et al. 2021). There is a possibility that MoT will spread to its neighboring and other countries in Asia as it spread from Brazil to other countries in South America. To prepare against its transmission to Japan, we initiated a breeding program for incorporating *Rmg8* into a Japanese elite cultivar. Consequently, we successfully produced its near-isogenic line (NIL) carrying *Rmg8*. A process of its breeding and its reactions to MoT in comparison with a 2NS-translocation line are reported here.

## MATERIAL AND METHODS

### Fungal material

We used five field isolates of *Pyricularia oryzae Triticum* pathotype (MoT), i.e., Br48, Br5 and Br116.5 collected in Brazil, BTMP-2(b) and BTGP-6(e) (abbreviated as T-105 and T-109, respectively, hereafter) collected in Bangladesh, and four transformants derived from Br48 (Table 1). Br48, T-105, and T-109 carry the eI type of *AVR-Rmg8* while Br5 and Br116.5 carry the eII and eII’ types, respectively. Br48ΔAVR-Rmg8_d6 (Br48ΔA8) is an *AVR-Rmg8* disruptant of Br48 produced by Wang et al. (2018) while Br48ΔA8+eI-3 is its transformant carrying the eI type of *AVR-Rmg8* derived from Br48 (Horo et al. 2020). Br48+PWT3 (M-16) and Br48+PWT4 (XB6) are Br48 transformants carrying *PWT3* and *PWT4*, respectively (Inoue et al. 2017).

**TABLE 1.**
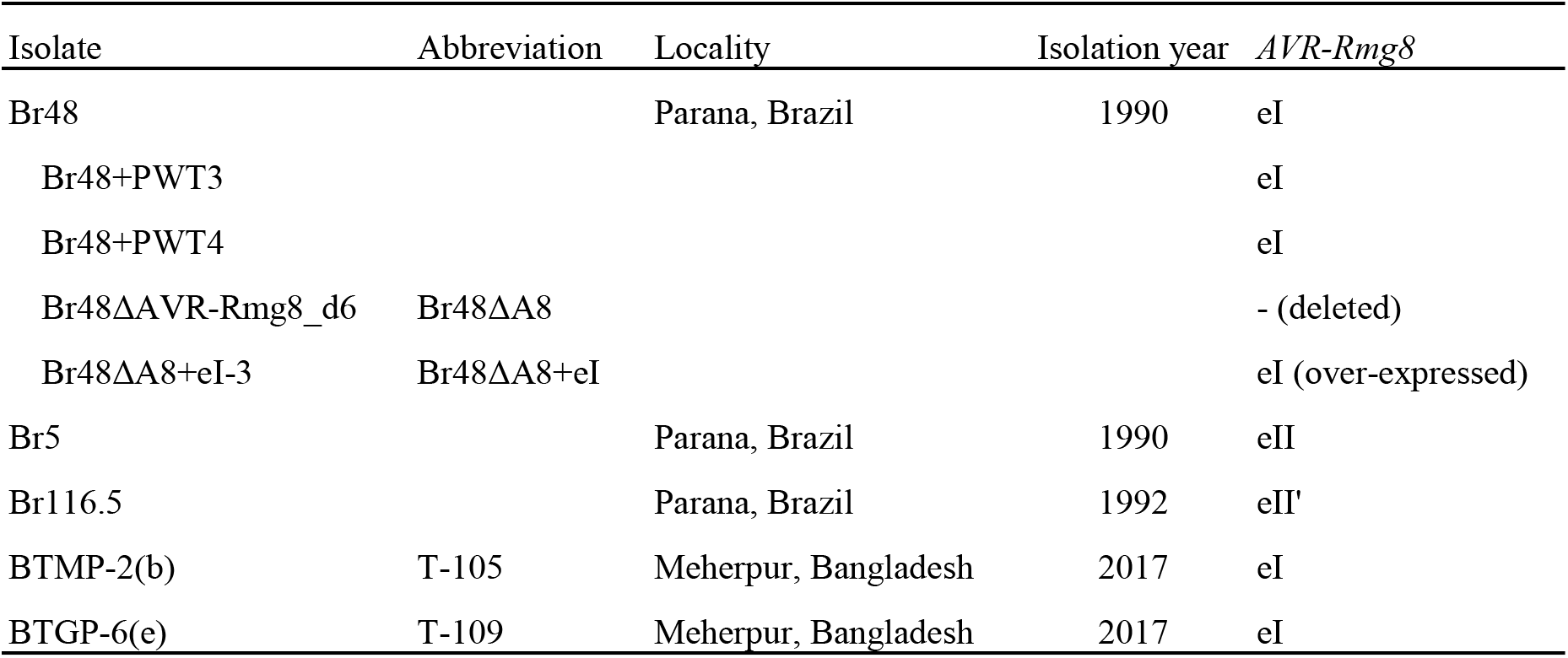
MoT isolates of *Pyricularia oryzae* and their derivatives used in this study

### Plant materials

Six Japanese elite cultivars of common wheat (*Triticum aestivum*) provided by National Agriculture and Food Research Organization (NARO) (Table 2) were first tested as candidates for a recipient of *Rmg8*. From these candidates ‘Chikugoizumi’ (ChI) was selected as the recipient (recurrent parent) on the basis of its genotype at the *Rwt4* locus (Inoue et al. 2021) and other characteristics. ‘S-615’ carrying *Rmg8* on the long arm of chromosome 2B (Anh et al. 2015) was employed as a donor of *Rmg8*. ‘Express’ and Express-2NS, a near-isogenic line (NIL) of ‘Express’ carrying the 2NS segment (Cruz et al. 2016; Helguera et al., 2003), were used for comparative analyses of *Rmg8* and 2NS. ‘Norin 4’ (N4) was used as a susceptible control.

**TABLE 2.** Genotypes of Japanese wheat cultivars at the *Rwt3* and *Rwt4* loci determined by inoculation with Br48 ormants carrying *PWT3* and *PWT4*

| Cultivar | Developer<sup>a</sup> | Year of registration | *Rwt3*<sup>b</sup> | *Rwt4*<sup>b</sup> |
| --- | --- | --- | --- | --- |
| Ayahikari | National Agriculture Research Center, Tsukuba * | 2000 | ND | ++ |
| Tamaizumi | NARO, Tsukuba | 2002 | + | ++ |
| Yumeshihou | NARO, Tsukuba | 2010 | + | - |
| Chikugoizumi (ChI) | Kyushu Agricultural Experiment Station, Chikugo * | 1994 | ++ | ++ |
| Shirogane-komugi | Kyushu Agricultural Experiment Station, Chikugo * | 1974 | + | ++ |
| Minaminokaori | NARO, Tsukuba | 2006 | - | ++ |

<sup>a</sup> NARO, the National Agriculture and Food Research Organization. Asterisked institutes have also been affiliated to NARO.
<sup>b</sup> ++, present; +, probably present; -, absent. ND, not determined due to an intermediate reaction.

### Seedling infection assay

Wheat seeds were germinated on wet filter paper for 24 hours, sown on soil (Sakata Prime Mix, Sakata Seed Corporation, Yokohama, Japan) in a plastic seedling case, and grown in a controlled-environment room with a 12-h photoperiod of fluorescent lighting at 22°C. Nine days after germination, primary leaves were fixed onto a hard-plastic board using rubber bands immediately prior to inoculation.

Fungal isolates/strains were grown on oatmeal agar media (made from 40 grams of oatmeal per liter and containing 2% of agar) for seven days. Aerial mycelia on the seven-day-old cultures were removed by swabbing the mycelial surfaces with cotton balls. The cultures were incubated further under blue light at 24–25°C for three to four days to induce sporulation. Conidia were harvested by scratching the surfaces of the cultures with microtubes to prepare a conidial suspension (1×10^5^ conidia/ml) containing 100 ppm of Tween 20. The suspension was evenly sprayed onto the fixed leaves using an air compressor. The inoculated leaves were placed in a humid tray sealed with plastic wrap, incubated in the dark at 22°C or 28°C for 24 hours, and then transferred to dry conditions with the same temperature and 12-h fluorescent lighting. Symptoms of the primary leaves were evaluated from 3 to 4 days after inoculation. Infection types were determined based on the size and colors of the lesions. The size was rated on six progressive grades from 0 to 5: 0 = no visible infection; 1 = pinhead spots; 2 = small lesions (< 1.5 mm); 3 = scattered lesions of intermediate size (< 3 mm); 4 = large lesion; and 5 = complete shriveling of leaf blades. A disease score (infection type) was represented with a number denoting the lesion size accompanied by a letter indicating the lesion color, B (brown) and G (green), as defined in Hyon et al. (2012). Infection types from 0 to 3 with brown lesions were considered resistant because the brown color was associated with hypersensitive cell death in this condition (Hyon et al., 2012). Infection types from 3G to 5G were considered susceptible. Seedling infection assays with intact primary leaves were conducted twice.

In the selection process for the NIL development, detached primary leaves were used for inoculation. Seedlings were grown in soil (Sakata Prime Mix) in a greenhouse (18– 26°C) for nine days. Primary leaves of the seedlings were cut out, inserted into Eppendorf tubes (2 ml) containing distilled water, and fixed onto a hard-plastic board. They were sequentially inoculated with Br48ΔA8 and Br48 as described by Nga et al. (2012). First, Br48ΔA8 was sprayed on the leaves whose bottom half was covered with aluminum foil, and incubated at 22°C for 24 h in a dark, humid box. Second, Br48 was sprayed on the same leaves whose upper half was covered with aluminum foil, and incubated in the dark, humid box for additional 24 h. Then, they were transferred to the dry condition with a 12-h photoperiod and incubated further at 22°C. Infection types were evaluated 3–4 days after the second inoculation. Leaves resistant to Br48 but susceptible to Br48ΔA8 were determined to be *Rmg8* carriers. Seedlings in the greenhouse that provided the *Rmg8*-carrying leaves were grown further to proceed to the next generation.

### Spike infection assay

Test plants were grown at the field in Kobe University, Kobe, Japan, for 4–5 months in the 2021–2022 season. At the stage between full head emergence and anthesis, the culms were cut and trimmed to 30–40 cm, and put into test tubes with water containing antibiotics and nutrition (MISAKI solution for cut flowers, OAT Agrio, Tokyo, Japan). The spikes, as well as their flag leaves, were inoculated with conidial suspension (1×10^5^ conidia/ml) containing 100 ppm of Tween 20 using an air compressor until their surfaces were evenly covered with fine droplets. The inoculated samples were sealed with a plastic bag, incubated in the dark at 25°C for 24 hours, and transferred to a dry condition at 25°C with 12-h fluorescent lighting. From five to nine days after inoculation, the infection type of the spike was rated with six progressive grades from 0 to 5: 0 = no visible infection; 1 = brown pinhead spots; 2 = brown lesions of intermediate size (< 3 mm); 3 = large lesions surrounded by brown tissues; 4 = extensive necrosis or chlorosis; and 5 = complete blighting of the spike. Infection types from 0 to 3 were considered resistant, and those from 4 to 5 were considered susceptible. The infection assay with spikes was conducted twice.

### Whole genome genotyping with GRAS-Di

GRAS-Di (Genotyping by Random Amplicon Sequencing-direct) is a whole genome genotyping technology developed by Toyota Motor Corporation (Enoki and Takeuchi, 2018). We genotyped eight BC_4_F_5_ individuals of #2-1-10 (*Rmg8*) and #4-2-10 (*rmg8*), respectively, with the GRAS-Di technology. Three individuals of ‘S-615’ and three individuals of ChI were also analyzed as parental lines. First, total genomic DNA was extracted from the seedlings following the published procedure (Yabe et al., 2014) with minor modifications. The quality of extracted DNA was confirmed with agarose gel electrophoresis. GRAS-Di library was prepared with a set of 12 random primers and sequenced with NovaSeq 6000 (paired-end, 150bp). Library preparation and sequencing were conducted by Eurofin Genomics K.K. Japan (https://eurofinsgenomics.jp/).

The generated fastq files were used for variant calling. First, the remaining adapters, low-quality reads, and unpaired reads were removed with fastp (v0.23.2). Next, trimmed reads were aligned to the v2.1 reference genome sequence of ‘Chinese Spring’ (IWGSC, 2018) with BWA (v0.7.8) using default options. Sequence variants were called from MAPQ > 40 reads with bcftools (v1.15.1) mpileup function. Subsequently, polymorphic sites fulfilling the following conditions were selected; QUAL value is larger than 100 and two or three samples of each parent have an opposite genotype call between ChI and S-615. Finally, the genotypes of the BC_4_F_4_ generation of the NILs #2-1-10 and #4-2-10 were estimated retrospectively from the genotypes of the eight individuals in the BC_4_F_5_ generation. The resulting set of polymorphic sites was used to draw the genomic landscape in the NIL lines produced through backcrossing.

## RESULTS

### Selection of *Rwt3* and *Rwt4* carriers from Japanese elite wheat cultivars

Wheat lines to be released to farmer’s fields are advised to carry *Rwt3* and *Rwt4* conditioning the resistance at the pathotype – plant genus level (Inoue et al. 2017). *Rwt3* and *Rwt4* recognize their corresponding avirulence genes, *PWT3* and *PWT4*, respectively, and therefore, their presence/absence can be inferred from comparisons of reactions to Br48 (a wild MoT isolate), Br48+PWT3 (a transformant of Br48 carrying *PWT3*) and Br48+PWT4 (a transformant of Br48 carrying *PWT4*) (Inoue et al. 2017). For example, if a cultivar is susceptible to a Br48 but resistant to Br48+PWT3 and Br48+PWT4, we can conclude that the cultivar carries both *Rwt3* and *Rwt4*. We inoculated seedlings of six representative elite cultivars with Br48, Br48+PWT3, and Br48+PWT4, and found that three out of the six cultivars carry both *Rwt3* and *Rwt4* (Table 2). From the three cultivars, we finally chose ‘Chikugoizumi’ (ChI) as a recipient of *Rmg8*.

### Introduction of *Rmg8* into ChI through recurrent backcrosses

The entire process of introduction of *Rmg8* is illustrated in Fig. 1. The starting material was an F_3_ line with the *Rmg8*/*Rmg8* genotype derived from a cross between a traditional Japanese cultivar ‘Shin-chunaga’ (*rmg8*/*rmg8*) and ‘S-615’ (*Rmg8*/*Rmg8*). The F_3_ line was recurrently backcrossed with ChI. In each generation, BC_n_F_1_ plants heterozygous for *Rmg8* were selected by inoculation with Br48 and Br48ΔA8. A BC_4_F_1_ plant heterozygous for *Rmg8* was self-pollinated, then phenotypes of the resulting BC_4_F_2_ plants were determined by inoculation (Fig. 2). Finally, the fixation or absence of *Rmg8* was confirmed in the BC_4_F_3_ generation. Two NILs in the BC_4_F_5_ generation (#2-1-10 with the *Rmg8*/*Rmg8* genotype and #4-2-10 with the *rmg8*/*rmg8* genotype) (Fig. 1) were used in the following assays.

**Fig. 1.**
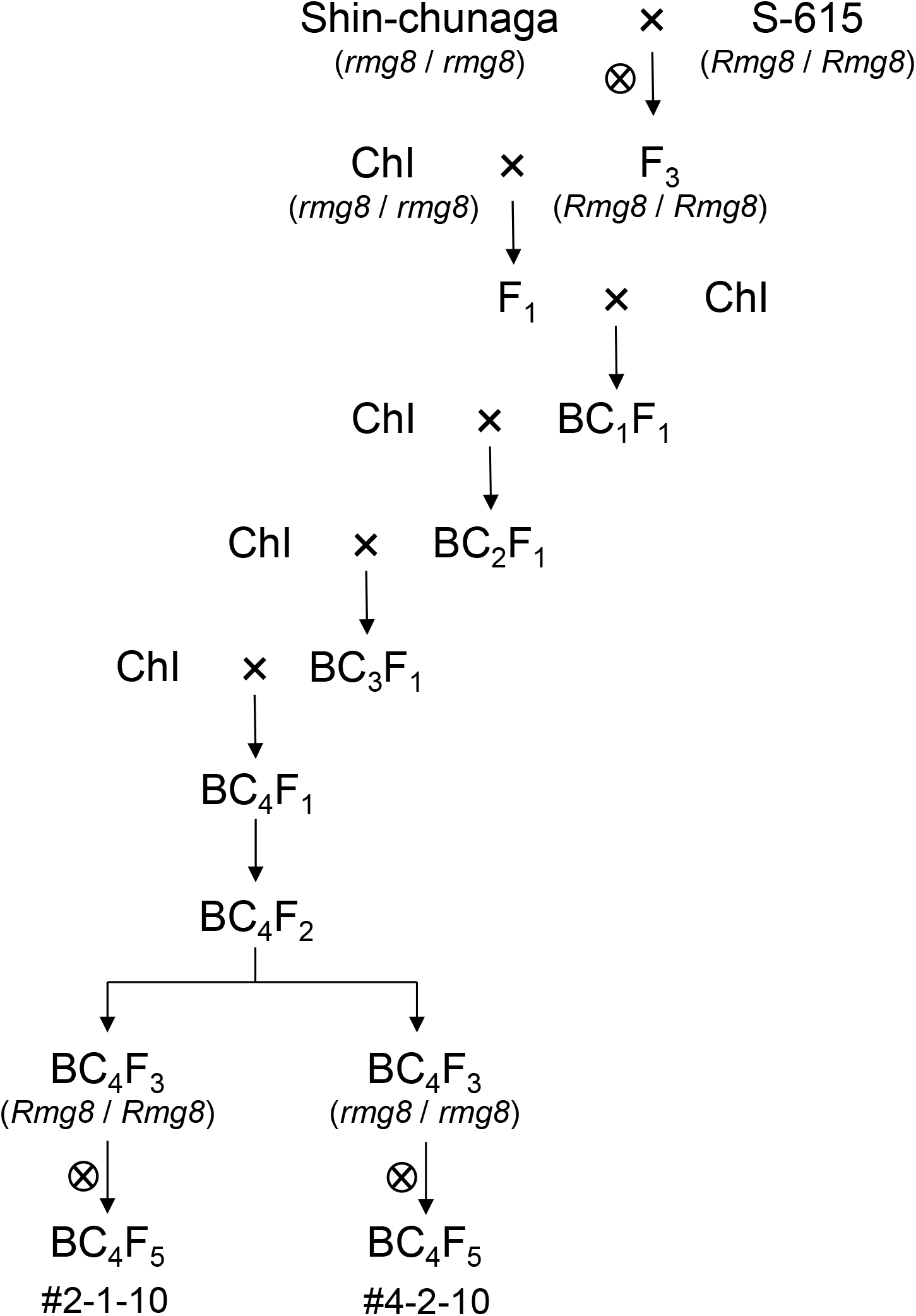
Flow diagram of the introduction of *Rmg8* from S-615 into Chikugoizumi (ChI).

**Fig. 2.**
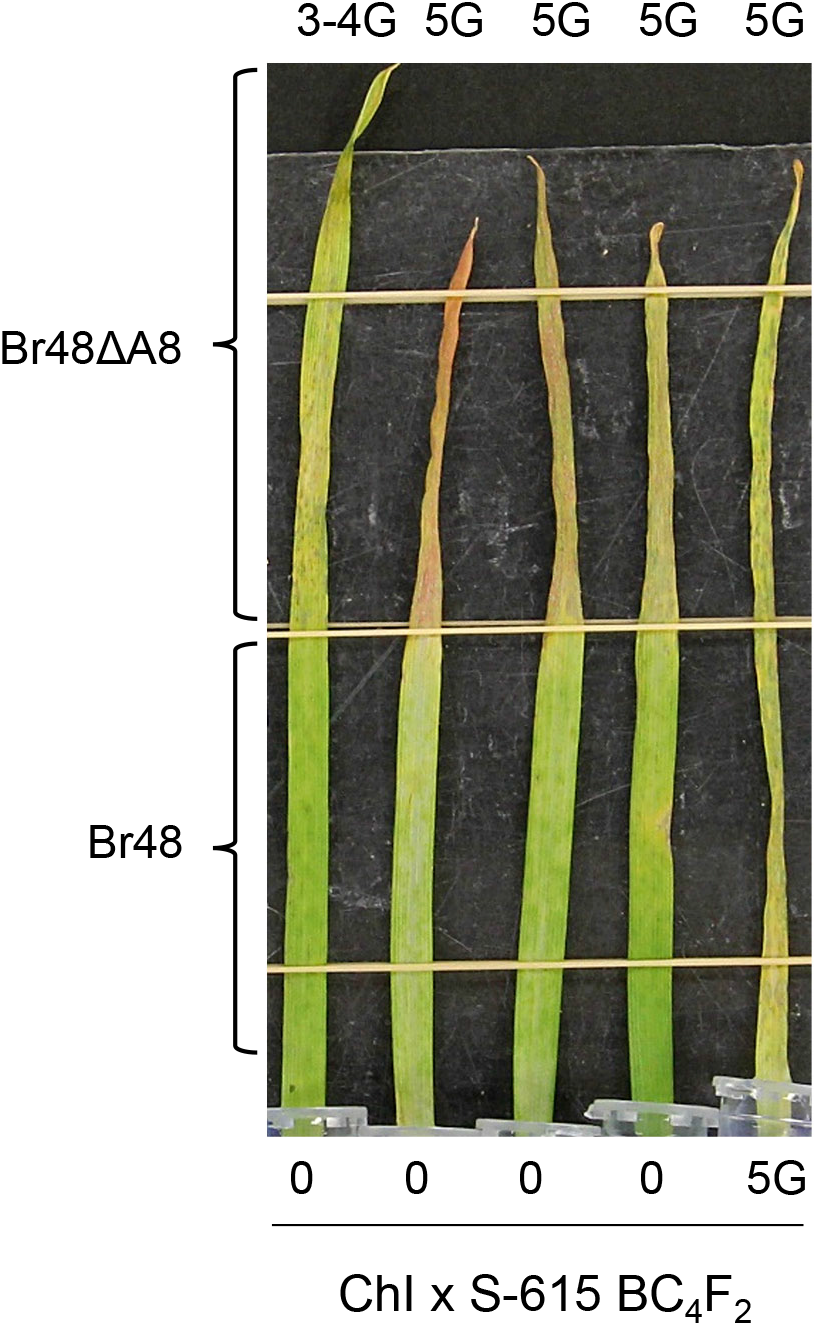
Symptoms on primary leaves of ChI x S-615 BC_4_F_2_ seedlings that were sequentially inoculated with two *P. oryzae* strains, 4 days after the second inoculation. The top half was first inoculated with Br48ΔA8 and incubated for 24 h in dark. Then, the bottom half was inoculated with Br48. Infection types with Br48ΔA8 and Br48 are shown at the top and bottom of the panel, respectively.

### Reactions of the NILs at the seedling stage

To evaluate the resistance of the developed NILs, their intact primary leaves were inoculated with representative wild isolates collected in Brazil (Br48, Br5, Br116.5) and Bangladesh (T-105, T-109), as well as Br48ΔA8 and Br48ΔA8+eI, and incubated at 22°C and 28°C. At 22°C, #2-1-10 (*Rmg8*/*Rmg8*) showed strong resistance with infection types 0–1B to all the wild isolates (Fig. 3). This resistance was compromised by the disruption of *AVR-Rmg8* (as shown against Br48ΔA8) but recovered by the reintroduction of the eI type of *AVR-Rmg8* (as shown against Br48ΔA8+eI) as expected. In contrast, ChI and #4-2-10 (*rmg8*/*rmg8*) were basically susceptible to all the isolates and transformants with infection types 4G–5G (Fig. 3). When observed carefully, however, ChI showed a slight degree of resistance to the Brazilian isolates, which #4-2-10 also inherited from ChI. This slight resistance was not recognized against the Bangladeshi isolates (T-105 and T-109); they caused complete shriveling of leaves of the *rmg8*/*rmg8* lines even at 22°C, suggesting that the Bangladeshi isolates are highly virulent (Fig. 3).

**Fig. 3.**
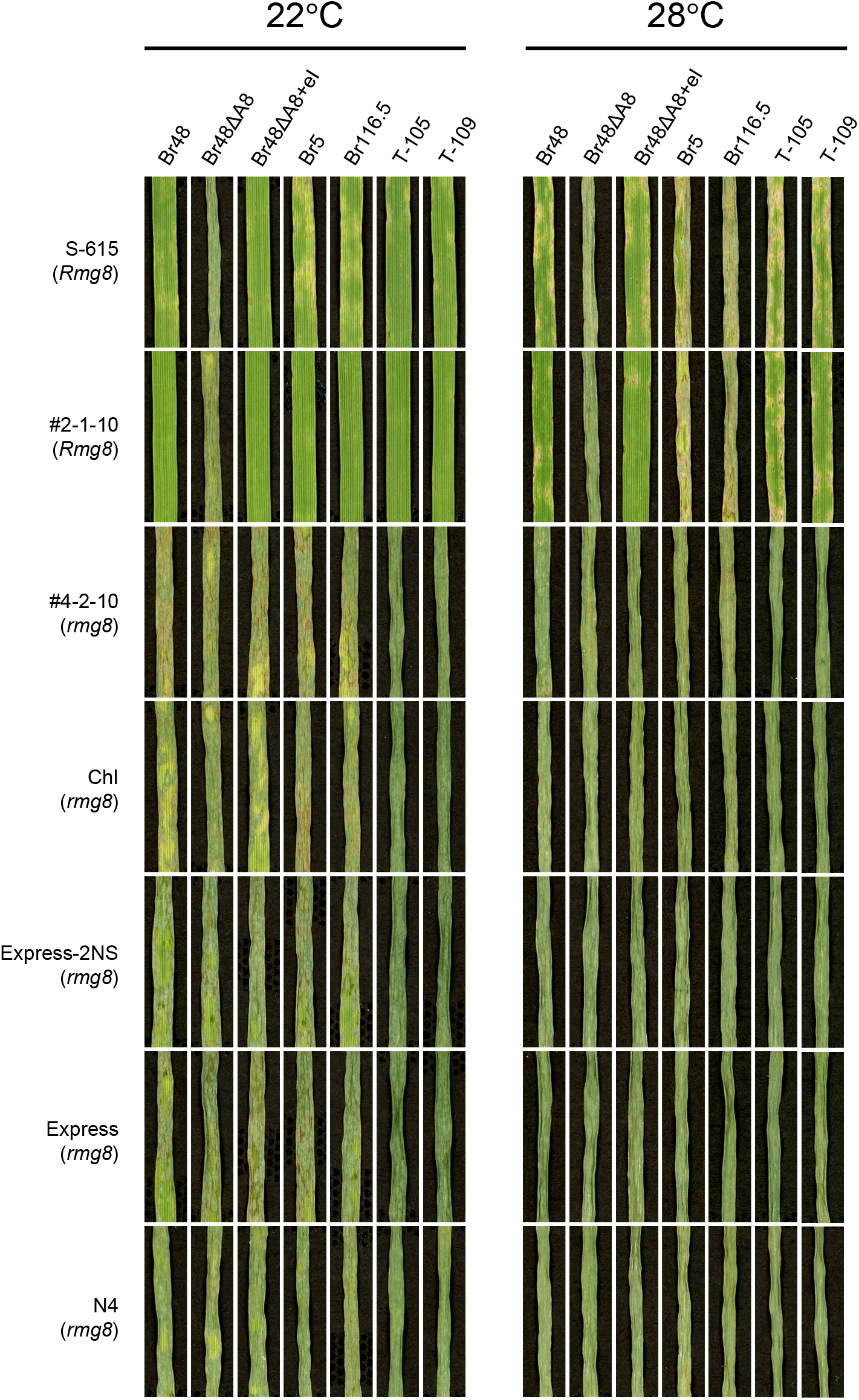
Reactions of intact primary leaves of wheat lines to MoT isolates collected in Brazil (Br48, Br5, Br116.5) and in Bangladesh (T-105, T-109) at 22°C (left) and 28°C (right). Br48ΔA8 and Br48ΔA8+eI are an *AVR-Rmg8* disruptant derived from Br48 and a transformant of Br48ΔA8 carrying the eI type of *AVR-Rmg8*, respectively. S-615, the donor of *Rmg8*; #2-1-10 and #4-2-10, the near-isogenic lines of cv. Chikugoizumi (ChI) carrying *Rmg8* and *rmg8*, respectively; Express-2NS, the near-isogenic line of cv. Express carrying the 2NS chromosomal segment; N4, cv. Norin 4 as a susceptible control.

At 28°C, #2-1-10 was still resistant to Br48, T-105 and T-109 (eI type carriers), but its resistance to Br5 and Br116.5 (eII/eII’ carriers) was compromised. Br5 and Br116.5 caused almost complete leaf shriveling in #2-1-10 and S-615 with slight tissue browning at 28°C. These results suggest that the interaction between *Rmg8* and eII/eII’ types of *AVR-Rmg8* is temperature-sensitive, and therefore, the eII/eII’ carriers are able to almost overcome the *Rmg8* resistance at higher temperature.

It should be noted that #2-1-10 exhibited stronger resistance than ‘S-615’ (the donor of *Rmg8*) against the wild isolates at 22°C. Express-2NS, the NIL of ‘Express’ carrying the 2NS segment, was susceptible to all the tested isolates at both temperature conditions, indicating that the 2NS segment is not effective at the seedling stage.

### Reactions of the NILs at the heading stage

To confirm the effectiveness of the developed NILs at the heading stage, their spikes and flag leaves were inoculated with Br48, Br48ΔA8 and Br48ΔA8+eI. Spikes of Express-2NS were resistant to all the three fungal strains at an equivalent level while those of ‘Express’ were susceptible, indicating that the resistance in spikes conferred by the 2NS segment is independent of *AVR-Rmg8*. However, flag leaves of Express-2NS were completely shriveled with all the three strains, indicating that the 2NS resistance is not expressed in flag leaves.

Spikes of #2-1-10 was resistant to Br48, susceptible to Br48ΔA8, and again resistant to Br48ΔA8+eI while ChI and #4-2-10 were susceptible to all the three strains (Fig. 4), confirming that the resistance of #2-1-10 in spikes is conditioned by *Rmg8*. The level of the spike resistance of #2-1-10 against Br48 was equivalent to that of Express-2NS. It should be noted that flag leaves of #2-1-10 showed the same pattern of reactions against the three fungal strains (Fig. 4), suggesting that *Rmg8* is effective not only in spikes but also in flag leaves in contrast to the 2NS segment.

**Fig. 4.**
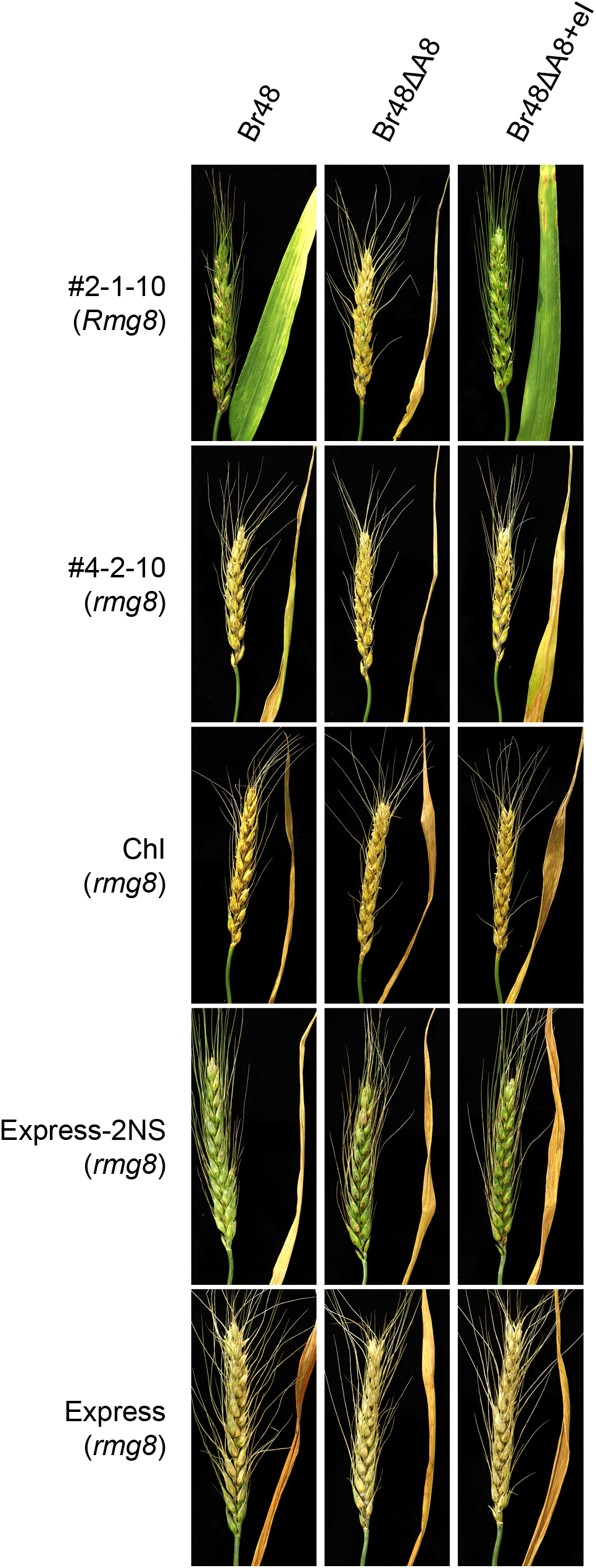
Reactions of detached wheat spikes and flag leaves to MoT isolate Br48, an *AVR-Rmg8* disruptant derived from Br48 (Br48ΔA8), and a transformant of Br48ΔA8 carrying the eI type of *AVR-Rmg8* (Br48ΔA8+eI) at 25 °C. #2-1-10 and #4-2-10, the near-isogenic lines of cv. Chikugoizumi (ChI) carrying *Rmg8* and *rmg8*, respectively; Express-2NS, the near-isogenic line of cv. Express carrying the 2NS chromosomal segment. Marker positions on chromosome 2B (Mbp)

### Estimation of genomic constitution of NILs

We performed whole-genome genotyping of #2-1-10 (a NIL carrying *Rmg8*), #4-2-10 (a NIL carrying *rmg8*), ‘S-615’ (the *Rmg8* donor), and ChI (the recipient) using the GRAS-Di technology to estimate how large portion of the ‘S-615’ genome was substituted with the ChI genome through recurrent backcrossing. Filtering of potentially polymorphic sites yielded a total of 15,215 variants (14,709 SNPs and 506 short indels) between ‘S-615’ and ChI across the 21 chromosomes. A comparative analysis suggested that at least 96.6% of the #2-1-10 genome was substituted with the ChI genome. On its chromosome 2B, an approximately 26.0 – 29.4 Mb region from the end of the long arm was derived from ‘S-615’ (Fig. 5), which was consistent with the mapped position of *Rmg8* reported in a previous study (Anh et al. 2015).

**Fig. 5.**
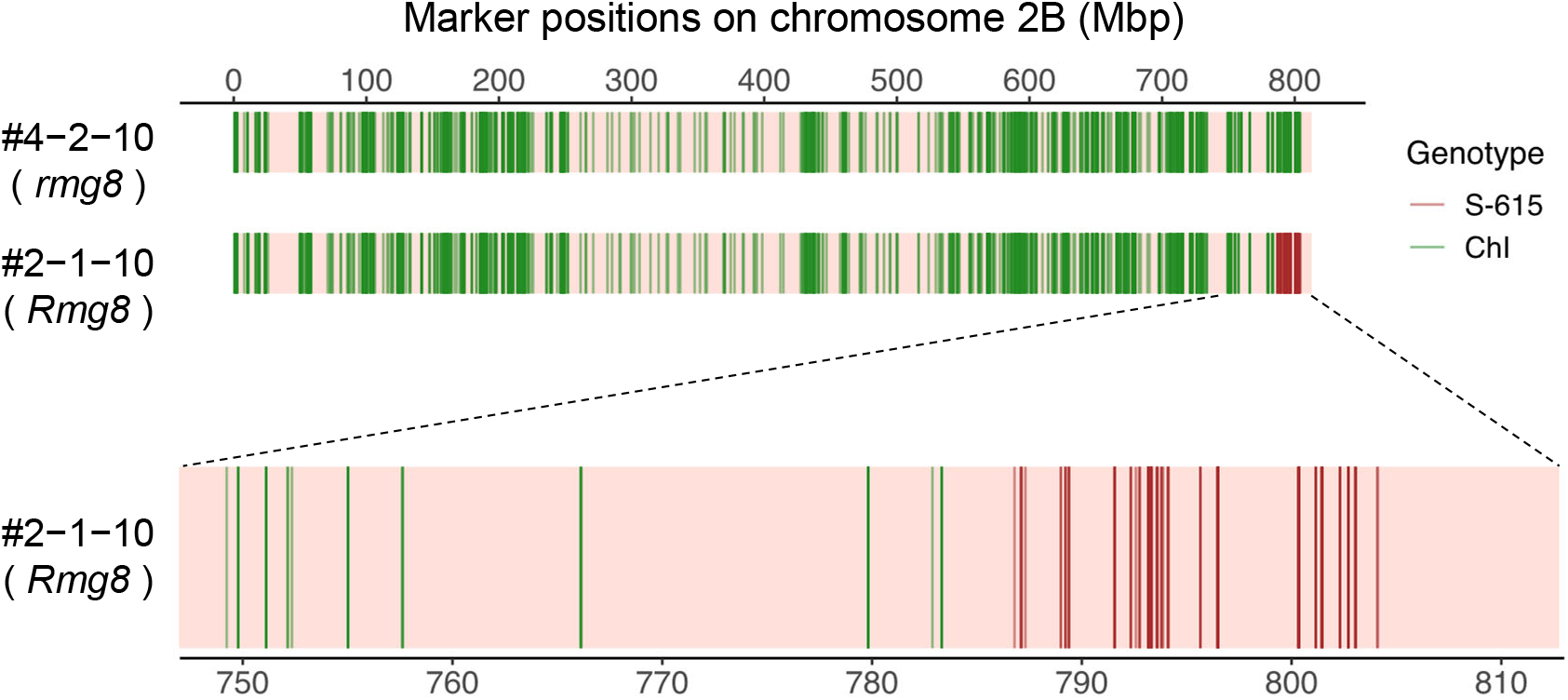
Graphical genotypes of chromosome 2B of the near-isogenic lines of cv. Chikugoizumi (ChI) carrying *Rmg8* (#2-1-10) and *rmg8* (#4-2-10) estimated by GRAS-Di.

## DISCUSSION

*P. oryzae* is composed of several host genus-adapted pathotypes such as the *Triticum* pathotype (MoT), *Lolium* pathotype (MoL), and *Eleusine* pathotype (MoE) which are pathogenic on *Triticum* spp., *Lolium* spp., and *Eleusine* spp., respectively (Valent et al. 2021; Kato et al. 2000; Tosa et al. 2004). Each pathotype forms a distinct lineage in a phylogenetic tree of *P. oryzae* isolates (Gladieux et al. 2018), but the *Lolium* lineage includes a unique isolate, Br58, which is specifically pathogenic on *Avena* spp. (Gladieux et al. 2018; Oh et al. 2002). *Rwt3* is a gene involved in the pathotype – genus specificity and conditions the resistance of common wheat to MoE, MoL, and Br58 (Asuke et al. 2020; Inoue et al. 2017). Wheat cultivars lacking *Rwt3* is considered to have served as a springboard for the host jump of MoL or its relatives to wheat, resulting in the emergence of MoT (Inoue et al. 2017). To prevent the recurrence of such host jumps, newly bred cultivars should carry *Rwt3*. *Rwt4* is another gene involved in the pathotype – genus specificity and conditions the resistance of common wheat to Br58 (Inoue et al. 2017). Inoue et al. (2021) reported that the *Rmg8*-mediated resistance was suppressed by *PWT4* but that the suppression was counteracted by *Rwt4*. This result suggests that newly bred cultivars whose resistance is conferred by *Rmg8* should also carry *Rwt4*. ChI fulfilled these two requirements. In addition, ChI is now widely deployed in southwestern district in Japan (a hot area with blast-conducive weather), and has a high dough quality for noodle production. Based on these considerations, we finally chose ChI as a recipient of *Rmg8*.

One of objectives of the present study was to compare the resistance conferred by *Rmg8* with the 2NS resistance. We had obtained six pairs of NILs, i.e., ‘Yecora Rojo’, UC1037, ‘Express’, UC1041, ‘Kern’, ‘Anza’, and their NILs carrying the 2NS segment (Helguera et al. 2003) from Dr. B. Valent, Kansas State University. We chose the Express/Express-2NS pair as a representative because our preliminary test with Br48 showed that the effect of 2NS at the heading stage was the most prominent in this pair. Cruz et al. (2016) reported that 2NS caused a significant reduction of spike blast but had no effect on leaf blast at the fourth leaf stage. However, it was not clear whether this was due to the difference of growth stages (the heading or seedling stages) or the difference of plant organs (spike or leaf). Our results showed that 2NS was effective in spikes but ineffective in primary and flag leaves (Figs. 3, 4). These results suggest that expression of the 2NS resistance is dependent not on growth stages but on plant organs. Further studies are needed to elucidate why the 2NS resistance is expressed only in spikes.

Wheat blast has been known to be mainly a spike disease in the field. Recently, however, leaf blast has been increasingly reported with epidemiological data suggesting its importance as a source of inoculum for spike blast (Valent et al. 2021). Based on these observations, Valent et al. (2021) suggested that resistance genes effective in the leaf stage should be introduced into varieties with head blast resistance. We suggest that, in the breeding program from now on, breeders should aim at producing wheat lines that express wheat blast resistance at both the seedling and heading stages, or more strictly in both leaves and spikes. Our results suggest that *Rmg8* confers resistance not only in leaves at the seedling stage (Fig. 3) but also in flag leaves and spikes at the heading stage (Fig. 4). Therefore, *Rmg8* will be useful to prevent the spread of MoT in Asia and Africa where the eI type carriers are prevailing. However, there is a high possibility that MoT in those continents will also overcome the *Rmg8* resistance (Jiang et al. 2021). Actually, isolates virulent or moderately virulent on *Rmg8* are already distributed in South America (Horo et al. 2020; Navia-Urrutia et al. 2022). A new breeding program for combining *Rmg8* with the 2NS segment is now under way.

## ACKNOWLEDGEMENTS

We express our gratitude to Jorge Dubcovsky, University of California Davis, U.S.A., for giving permission for us to use the 2NS NIL lines, and Barbara Valent, Kansas State University, U.S.A., for providing seeds of the NIL lines, useful information on them, and valuable comments on the manuscript. We also thank Md Tofazzal Islam, Bangabandhu Sheikh Mujibur Rahman Agricultural University, Bangladesh, for providing the wheat blast isolates collected in Bangladesh. Computations were partially performed on the NIG supercomputer owned by National Institute of Genetics, Research Organization of Information and Systems. This research was supported by the research program on development of innovative technology grants (JPJ007097) from the project of the Bio-oriented Technology Research Advancement Institution (BRAIN).

